# Multi-omic characterization of the thermal stress phenome in the stony coral *Montipora capitata*

**DOI:** 10.1101/2021.02.05.429981

**Authors:** Jananan S. Pathmanathan, Amanda Williams, Timothy G. Stephens, Xiaoyang Su, Eric N. Chiles, Dennis Conetta, Hollie M. Putnam, Debashish Bhattacharya

## Abstract

We used network methods to analyze transcriptomic and polar metabolomic data generated from the stress resistant stony coral *Montipora capitata*. Corals were exposed to ambient or thermal stress conditions over a five-week period that coincided with a mass spawning event of this species. Gene co-expression networks showed that the early thermal stress response involves downregulation of growth and DNA replication, whereas signaling and the immune response are strongly upregulated. Later stages are dominated by suppression of metabolite transport and biomineralization and enhanced expression of transcriptional regulators. Integration of gene-metabolite data demonstrates that the major outcome of the thermal treatment is activation of animal redox stress pathways with detoxification of reactive oxygen species being dominant. Differential regulation of the highly conserved cytochrome P450 gene family was of particular interest with downregulation of CYP1A1, involved in progesterone metabolism, potentially explaining the attenuated mass spawning observed during the sampling period.

## Introduction

Coral reefs are vitally important natural resources because they are home to about one-quarter of all marine biodiversity (1). Since their radiation in the Middle Triassic period ~ 240 million years ago (Ma) (2), stony corals have survived five mass extinction events (3). Their long-term survival underscores the inherent resilience of these holobionts, that is comprised of the cnidarian host, algal symbionts, prokaryotic microbiome, and viruses. But does past performance predict future success, given the unfolding Anthropocene extinction event (4)? Environmental shifts can lead to destabilization of the symbiosis (dysbiosis) between the coral animal and its partners (5), resulting in mass bleaching events and misregulation of spawning (6).

Given these concerns, we undertook a study of the coral thermal stress phenome to identify key genes and metabolites (both known and “dark”, of unknown function) that sustain the holobiont under heat stress. Integration of gene-metabolite data was used to identify the major pathways of stress response that form targets for intervention to improve coral resilience. Our model system, *Montipora capitata* (Figure 1A), is a stress resistant, hermaphroditic, broadcast spawning Hawaiian coral that has the ability to sustain itself during periods of bleaching through heterotrophic feeding (7). *M. capitata* nubbins (coral fragments) were subjected to ambient (27°C) and thermal stress (ambient + ~3°C) conditions over a 5-week period, during which time multi-omics (transcriptomic and polar metabolomic) data were collected at three different time points from the same genotypes and analyzed using network approaches (8, 9). To study coral animal gene-holobiont metabolite interactions, we used the computer program Metabolite Annotation and Gene Integration [MAGI (10)] to link metabolite features to biological pathways that showed gene expression differences. The multi-omic integration results provide strong evidence of the coral animal response to redox stress, including the scavenging of excess molecular oxygen. We also find evidence that progesterone metabolism may play a role in misregulation of mass spawning due to thermal stress.

**Figure 1:**
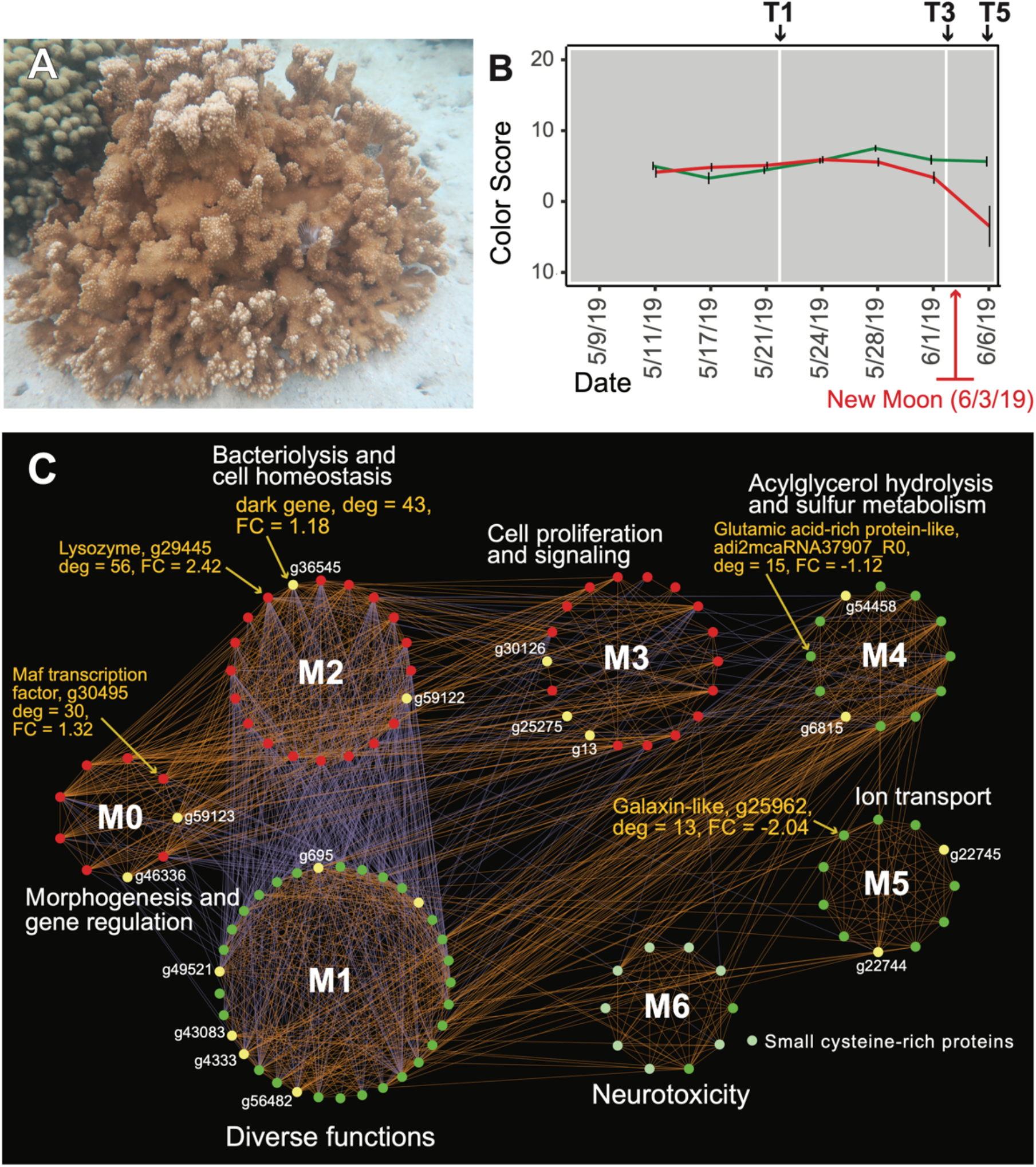
Analysis of the rice coral *Montipora capitata*. (A) *M. capitata* photographed in waters near the Hawaiʻi Institute of Marine Biology (HIMB) in O‘ahu, HI. Image taken by D. Bhattacharya. (B) Color scores and their standard errors for the ambient (green line) and high temperature (red line) treated *M. capitata* nubbins that were cultured in tanks at HIMB. Low color scores indicate the bleaching phenotype in coral holobionts. The omics data sampling points are shown with the white lines at T1 (5/22/19), T3 (6/03/19), and T5 (6/07/19) [for details, see (11)]. The date of the New Moon in June 2019 is also shown. (C) Network of differentially expressed genes in *M. capitata* at TP1 (the early thermal stress phenome) showing the different gene modules and their interactions. Red nodes are up-regulated and green nodes are down-regulated. The yellow nodes indicate unidentified, or dark genes with *M. capitata* gene IDS shown. Degree (number of edges linked to a node) and fold-change (FC) in gene expression, which are measures of gene importance in the stress response, are shown for one dark and four annotated genes. The down-regulated genes in M6 that is dominated by members of the small cysteine-rich protein family, often involved in signaling and protein interactions, are shown with the light green nodes. The module annotations are a general representation of overall function(s) (see Supplementary File 1 [TP1 full network] for annotation of each gene in the network).

## Results

### Culture conditions

In our experimental set-up, water was drawn from Kāne‘ohe Bay, O‘ahu and heated to 2.7° - 3.2°C above ambient temperature in tanks at the Hawaiʻi Institute of Marine Biology [for details, see (11)]. Color scores, which are a proxy for algal abundance and coral health [i.e., a low score indicates bleaching (12)] were recorded for the ambient and stress treated nubbins (Figure 1B). The period of sampling (late May to early June 2019) coincided with one of the three annual mass-spawning events for *M. capitata* in the region. Therefore, genes and metabolites involved in coral reproduction were expected to be present in the RNA-seq and polar metabolomic data. It should be noted that the eukaryotic RNA-seq data was filtered to only include reads that mapped to the predicted *M. capitata* protein-coding genes (13). Therefore, the gene expression results are limited to the coral animal, whereas the metabolomic data are derived from the entire holobiont. All omics data reported here were collected at time points T1, T3, and T5 (Figure 1B). The color scores demonstrate little impact until T3, followed by strong evidence of bleaching by T5. The network statistics reflect this temporal growth in complexity of the stress response (Figure 1-table supplement 1). Because there was an unexpected warming event in Kāne‘ohe Bay during the experiment that increased the ambient seawater temperature by ca. 2°C (11), we expected the gene co-expression data to show evidence of thermal stress at T1, that should become more pronounced at T3 and T5. In the networks of differentially expressed genes (DEGs) presented below (see Methods for details), we discuss different modules of the polygenic thermal stress response in *M. capitata*.

### Gene Co-Expression Network Analysis

#### The early stress response

Consistent with our prediction of early stress, inspection of the full network of DEGs, shows stress pathways [Module 2, M2; Figure 1C (Supplementary File 1, TP1 full; Cytoscape file with gene annotations] that are moderately upregulated at T1 and are negatively correlated with the expression of modules with a variety of different functions. In M1, down-regulated genes include a Mab-21 domain-containing protein involved in development (*M. capitata* gene g58637, fold-change, FC = −1.35), a tetraspanin involved in organizing plasma membrane protein-protein interactions (g57322 FC = −1.17) (14), and proliferating cell nuclear antigen involved in regulation of DNA replication and replication-associated processes (PCNA, g43600 FC = −1.09) (15). The T1 network also shows the strong up-regulation in M3 of genes involved in signaling (e.g., brichos domain-containing proteins; g29710 FC = 5.94, g29707 FC = 5.40) and a previously described protein in sea urchin and sea cucumber involved in embryo development [fibropellin-1 (16, 17), g71193 FC 5.47; see also Supplementary File 1]. Within M2 are well-characterized genes such as C-type lysozyme that is involved in bacteriolysis and the immune response (g29445 FC = 2.41) (18). Other genes in this module encode a PiggyBac transposable element-derived protein 5 (g9717 FC = 1.02), which is a potential driver of genome rearrangements and a putative growth factor receptor (TNFR-Cys domain-containing protein, g20162 FC = 1.12). Two genes with potential roles in biomineralization, a glutamic-acid rich protein and a galaxin-like gene, whose products are associated with the coral skeletal organic matrix (SOM) (19) are down-regulated in modules M4 and M5, respectively.

Also occupying key positions in the T1 network (Figure 1C) are genes of unknown function (“dark genes”) that are shown as yellow nodes with gene IDs indicated (Supplementary File 2). We define dark genes as predicted proteins with RNA-seq validation that are either lineage-specific or shared with other species and do not share significant sequence similarity (BLASTP *e*-value cut-off ≤ 1e^−5^ against the nonredundant NCBI database). Dark genes are either novel or too highly diverged to identify putative homologs in existing data, although some may encode a known domain associated with novel sequence (20). For example, 33% of dinoflagellate algal genes lack an annotation, but 1.4% of these unknown proteins contain a known domain (21). In the network at T1, we highlight *M. capitata* dark gene g36545 that has a weak hit to a N-terminal death-domain superfamily (*e*-value = 6.79e^−04^). This sequence is a member of a gene family that is upregulated in module M2 and has one of the highest degree values [number of edges linked to a node: i.e., these genes may act as regulatory components of the transcriptional network (13)] in this module (Figure 1C). Analysis of protein distribution demonstrates that dark gene g36545 is shared by, and limited, to other stony corals (Figure 1-figure supplement 1).

#### Downregulated genes in the later stress response

Next, we focused on the networks generated from the T3 and T5 DEG data for *M. capitata*. These networks are larger than the T1 network; each comprising 20 modules (Figure 2-figure supplement 1). We identified some genes with high degree in these networks, as well as dark genes, but will focus here on individual modules in the T5 network to gain insights into the later stage of the thermal stress response. M1 in the TP5 network (Figure 2) contains many significantly down-regulated genes that are dominated by metabolite transporters. These include a variety of sodium-coupled transporters shown with the magenta nodes, including a sodium-coupled neutral amino acid transporter (gene adi2mcaRNA35257_R0, deg = 3 FC = −1.61), a sodium-coupled monocarboxylate transporter 1 (g37389 FC = −1.53) putatively involved in the transport of a variety of substrates including short-chain fatty acids and lactate (22), and a probable sodium/potassium-transporting ATPase subunit (g39446 FC = −1.68) involved in the sodium-coupled active transport of nutrients (23). The transporter with the highest degree in this module (deg = 36) is a putative ammonium transporter (g26425, deg = 36, FC = −1.29; Figure 2). The suppression of metabolite transport by the coral host may potentially be a response to reduction in algal productivity. More likely, it indicates redox stress caused by algal symbionts that leads to the generation of reactive species due to dysfunction in electron transport [see below (24)]. The inhibition of organic carbon production by the algae, precipitated by prolonged thermal stress (25), can lead to their expulsion, resulting in bleaching (26). That is, in addition to a role in host processes, the coral animal may be dampening algal proliferation by reducing access to nutrients needed for growth such as ammonium, as demonstrated in the cnidarian model *Aiptasia* under the fully symbiotic stage (27). This hypothesis conflicts with the findings of Fernandes et al. (28) who found that ammonium enrichment reduced thermal stress in the coral *Stylophora pistillata* and supported symbiont stability. This aspect may be less important for Hawaiian *M. capitata* that meets 100% of its energy needs through heterotrophic feeding during periods of bleaching (7).

**Figure 2:**
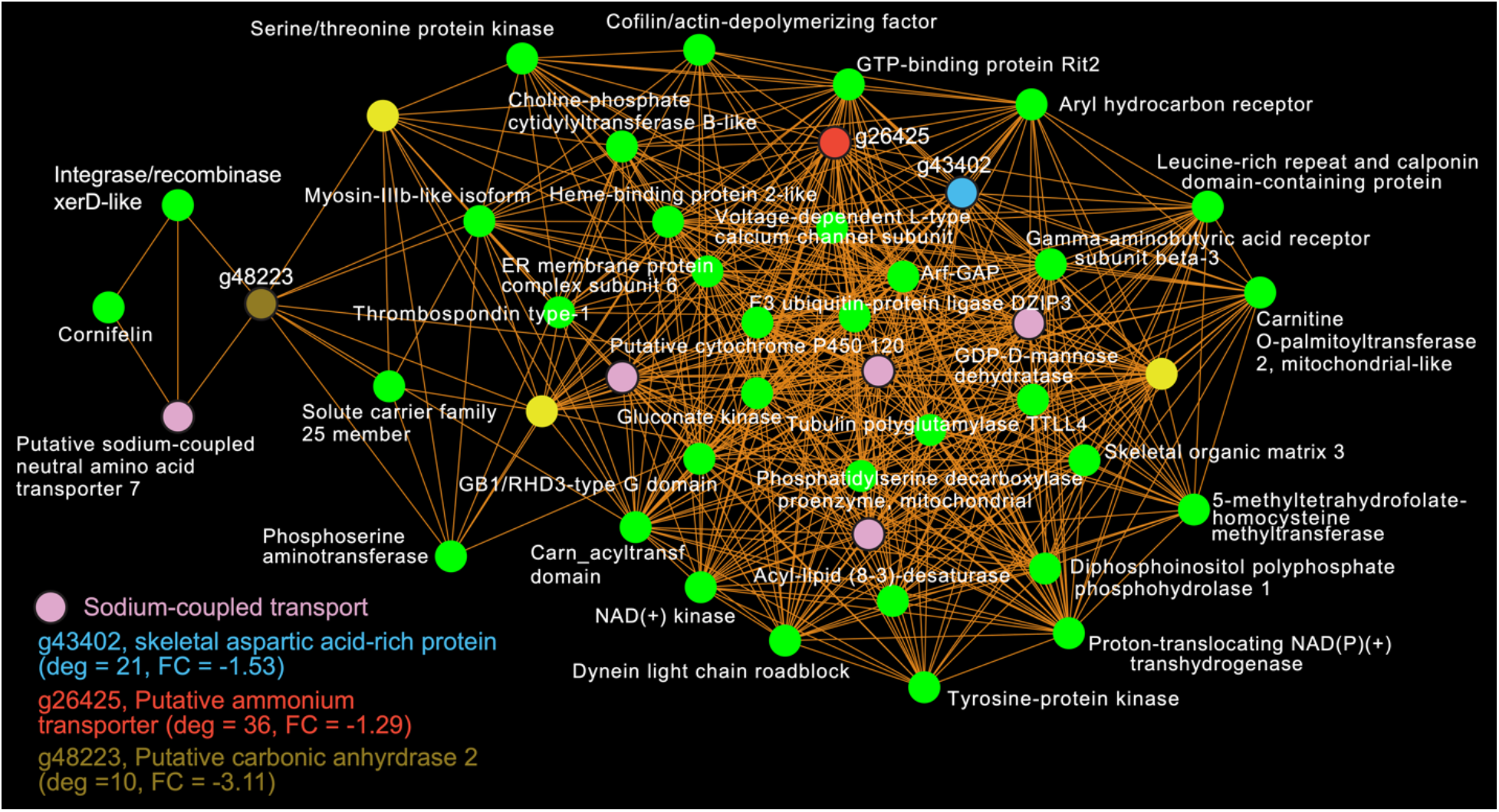
*M. capitata* TP5 module M1 comprising significantly down-regulated genes. Yellow nodes are dark genes and annotations are provided for three genes of interest (see legend). Degree, fold-change (FC), and gene annotation are shown for the genes discussed in the text. Genes encoding sodium-coupled transporters are shown with the pink nodes.

Another key component of module M1 is the skeletal aspartic acid-rich protein 3 that is related to coral acid-rich protein 4 (CARP4; ca. 40% protein identity) involved in biomineralization (CaCO_3_, aragonite in corals). CARPs are largely independently derived, secreted proteins rich in glutamic and aspartic acid residues that accumulate in the calicoblastic space of corals, playing roles in calcification (29–31). CARP-encoding genes are differentially expressed during coral development with CARP4 and CARP5 being strongly up-regulated in the calcifying spat stage of *Pocillopora damicornis* (32, 33). In M1, a single CARP is present, that is centrally located in the network (deg = 21) and strongly downregulated at TP5 (FC = −1.53). A maximum likelihood phylogeny of this protein (Figure 2-figure supplement 2) shows this gene to be present in non-coral species and to have undergone ancient gene duplications (provisionally named C1-C4 and R1-R4 for complex and robust coral species, respectively) within the scleractinian lineage with *M. capitata* encoding divergent paralogs. However, only the gene (g43402) encoding CARP4 is significantly downregulated under thermal stress in this species. These results indicate that the *M. capitata* thermal stress phenome includes suppression of the biomineralization reaction (also evident in TP1, see above) with the concomitant down-regulation of a putative carbonic anhydrase 2 (Figure 2), a zinc metalloenzyme that catalyzes the reversible hydration of carbon dioxide to bicarbonate (34). The large subnetwork in this module is connected to three peripheral genes, bridged by the putative carbonic anhydrase, of which one is the sodium-coupled neutral amino acid transporter and another is an integrase/recombinase (recombination function) xerD-like protein (g51825 FC = −1.64). A gene encoding an uncharacterized SOM protein 3 (g29122 FC = −1.02) is also downregulated under thermal stress.

#### Up-regulated genes at T5

Another module of interest in TP5 is M4 (Figure 3A), that contains a variety of significantly up-regulated genes with roles in signaling and immunity (e.g., netrin receptor UNC5C [g6679 FC = 2.00], two neuronal pentraxin-like genes [g46559, g46566 FC = 1.79, 2.45, respectively]) and transcriptional regulation (e.g., BTG1 protein [g32300 FC = 1.27], MafB [g30496 FC = 2.36], thyrotroph embryonic factor [g57753 FC = 1.42]). BTG family members are transcriptional regulators that can enhance or repress the activity of transcription factors. Maf proteins are widespread among metazoans, including corals, and are bZIP (basic [DNA-binding] and leucine zipper [homo- or heterodimerization] domains) transcriptional factors that are involved in oxidative stress and detoxification pathways (34, 35). Multiple Maf genes are upregulated at TP5 in M2, including *mafF* (g30493 FC = 2.39) and two Maf domain-containing proteins (g30494, g30495 FC = 1.96, 2.04, respectively). Two Maf domain-containing proteins are present in M18 (g2209, g26625 FC = 1.41, 1.24, respectively). In M4, the pentraxin domain-containing proteins are of interest because these are multimeric, calcium-binding proteins often involved in immunological responses (36). Located in this module are two proteins that interact with calcium: one is a calcium-binding EF-hand protein (g14108 FC = 1.67) and the second is a calcium-activated potassium channel subunit (g16479 FC = 1.86).

**Figure 3:**
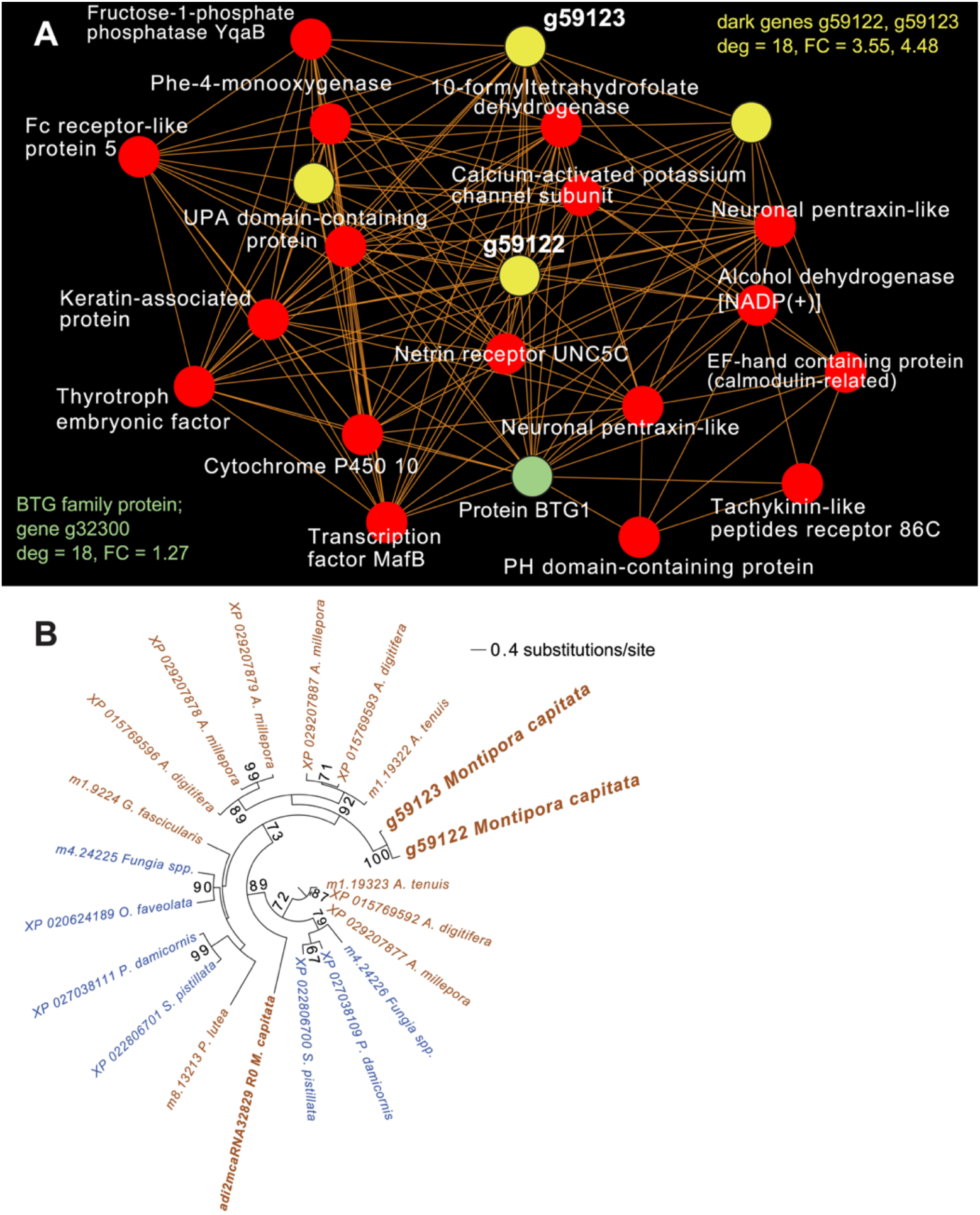
*M. capitata* TP5 module M4 comprising significantly up-regulated genes. Yellow nodes are dark genes. Degree and fold-change in gene expression are shown for the genes discussed in the text. (B) Maximum likelihood [IQ-Tree (66)] phylogenetic analysis of paralogous coral dark genes g59122 and g59133 and related homologs inferred using default parameters and 1000 ultrafast bootstrap replicates. The results of the bootstrap analysis are shown on the branches when >60%. The legend shows the expected substitution rate for the protein dataset. Complex and robust coral species are shown in brown and blue text, respectively.

Embedded within this network of conserved stress response proteins are 4 dark genes, two of which are paralogs that comprise highly connected hubs in this module (g59122 and g59123, both have deg = 18 and lack a domain hit using CDSEARCH). This gene family was only detected in stony corals (Figure 3B) and offers a promising target for functional analysis. Both of these dark genes show a high fold-change in gene expression when compared to ambient conditions (g59122, FC = 3.55) with g59123 having the highest value in this module (FC = 4.48).

### Gene-metabolite interactions

Many advances have been made in identifying individual gene and metabolite markers of coral thermal stress (5), but little has been done to link these two omics data sources. This is explained by the complexity of holobiont metabolomic interactions, combined with the massive number of dark genes and dark metabolites in corals for which currently no function, and therefore no causal relationship exists (11). Approaches such as MAGI, which link metabolites and gene expression data to known biological pathways provides a foundation for studying non-model systems (10). The MAGI approach was applied to the *M. capitata* polar metabolite and transcriptome datasets, using a stringent MAGI score ≥ 5 as the cut-off to ensure reliability (see Figure 4-table supplement 1 and Methods), to discover links between them that could clarify the coral thermal stress response.

#### Animal response to redox stress

Analysis of the MAGI output provided clear evidence of redox stress in the coral animal (Figure 4), with 21/27 of the high-confidence upregulated reactions having oxidoreductase functions (Figure 4-table supplement 1). Of these 21 reactions, 10 involve O_2_ as a substrate and the release of a water molecule, the majority of which include cytochrome P450 domains. The rate of metabolism at higher temperatures increases and can lead to physiological hyperoxia. Under elevated temperatures, oxygen absorbs excitation energy and becomes active in the form of superoxide radicals and hydrogen peroxide (37). These reactive oxygen species (ROS) are likely to be key contributors to coral thermal stress (38, 39).

**Figure 4:**
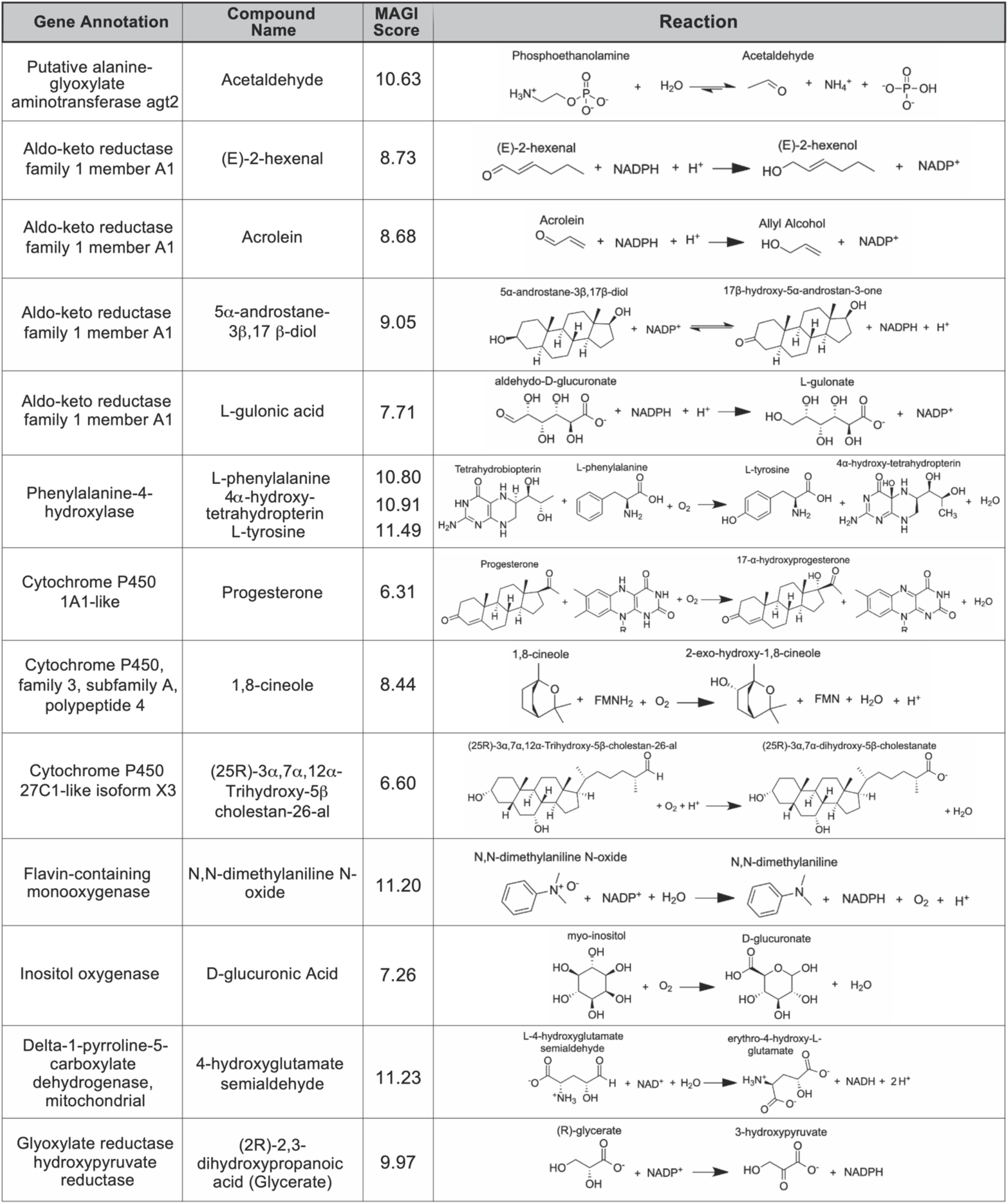
Results of the MAGI analysis. This image shows some of the pathways that are upregulated under thermal stress at TP5. MAGI scores are shown for each gene-metabolite pair.

#### Upregulation of the phenylalanine-4-hydroxylase pathway

A pathway of particular interest with regard to the coral thermal stress response involves phenylalanine-4-hydroxylase (P4H), which is a homotetramer of four phenylalanine hydroxylase (PH) enzymes, each containing three domains (a regulatory N-terminal domain, a catalytic domain, and a C-terminal domain) that use a non-heme Fe(II) cofactor (40). P4H catalyzes the bidirectional reaction of L-phenylalanine to L-tyrosine with (6*R*)-L-*erythro*-5,6,7,8-tetrahydrobiopterin (BH4) as a cofactor (Figure 4). The gene expression and metabolite integration results show upregulation of the *p4h* gene (FC = 1.27), as well as increased ion counts for all reaction participants except BH_4_. BH_4_ donates two electrons to reduce the iron atom to ferrous iron and cleaves O_2_ to reduce phenylalanine (Phe) to tyrosine (Tyr). Molecular oxygen can oxidize the ferrous iron, regenerating the enzyme. In this pathway, 4α-hydroxy-tetrahydropterin is first dehydrated and then reduced by an NADH-dependent component of P4H, the phenylalanine hydroxylase stimulator (PHS) (41). Phe and Tyr are both synthesized by scleractinian corals, either from intermediates in glycolysis, gluconeogenesis, the pentose phosphate pathway (PPP), the tricarboxylic acid cycle (TCA), or the pentose phosphate shunt, depending on the substrate used in previous studies (42). When coral samples are incubated with ^14^C lysine, Tyr and Phe are produced through gluconeogenesis, glycolysis, or the PPP following the TCA cycle. Corals can also take up dissolved free amino acids from the surrounding sea water (43). These results could explain the lack of reactant depletion in the P4H pathway. Although P4H can function bidirectionally, it is more likely that the enzyme is reducing Phe to Tyr. The reverse reaction is not energetically favored because P4H preferentially binds Phe rather than Tyr, and one of the most important biological roles of Phe is producing Tyr, a substrate for receptor tyrosine kinases that are implicated in the coral stress response (44).

#### Misregulation of spawning

Cytochrome P450-like (*CYP-like*) genes, which facilitate the biotransformation of important intracellular compounds (45), are implicated in four metabolic pathways in our results. The most notable is the downregulation of progesterone, putatively *via* a *CYP-like* gene (FC = −1.10), that coincides with the June 2019 *M. capitata* mass spawning event. Progesterone, a sex steroid, can be produced in multiple ways, but usually involves *β*-hydroxylation catalyzed by CYP enzymes (46). Many examples of CYP enzymes capable of metabolizing progesterone occur among metazoans, such as CYP17 dehydrogenase (CYP17) in scleractinian corals (47). Existing data suggest that sex steroids regulate scleractinian reproduction (48). CYP17 converts progesterone to androgens, and in the absence of thermal stress, the enzymatic activity of CYP17 remains constant over the lunar cycle in the brooding coral *P. damicornis* (49). Twan et al. (50) found that androgen production increased prior to spawning in the coral *Euphylia ancora*.

## Discussion

Coral reefs are under worldwide threat from warming oceans and locally, due to human activities such as over-fishing, the discharge of pollutants, and uncontrolled development (5). To address these pressing issues, we need robust and reliable approaches to identify thermal stress-related genes and pathways for genetic modification, or selection using controlled breeding to generate more resilient coral lines (5). Our study provides important advances in these areas that are described above. Two aspects of the results deserve further discussion.

The first is the gene-metabolite interaction analysis of the phenylalanine-4-hydroxylase pathway in which BH_4_ was unexpectedly absent in the MAGI results. Some plausible explanations for this result are as follows. In the kinetic model, P4H requires BH_4_, Phe, and O_2_ to be bound, in that order (51). BH_4_ binds first, converting the enzyme to its inactive form, E_i_, until sufficient Phe is present in plasma, at which point Phe binds and converts P4H to its activated form, E_a_ (52); BH_4_ bound to P4H would not have been detected in our analysis. Given that BH_4_ is involved in other cellular functions it is possible that its levels might be depleted under heat stress, despite upregulation of the P4H pathway. This is relevant when considering the stoichiometry of the reaction, specifically, the number of BH_4_ molecules needed as cofactors depends on cellular conditions. Higher pH and temperature may require more than one BH_4_ to reduce two iron atoms (53), further reducing the number of free BH_4_ molecules available for detection. It is also possible that another tetrahydropterin was used as an electron donor, which could result from two possible scenarios. The first is that NADH is required to regenerate BH_4_, which might promote the use of other compatible substrates when energy is limited. This would explain the accumulation of dihydrotetrapterin. A tetrahydropterin ring with a phenyl, ethyl, hydroxymethyl, or trihydroxypropyl substitution at the 6 carbon or a methyl group at the 7-carbon can sustain P4H hydroxylation (40). In addition, the entire pyrazine ring is not needed for P4H activity, therefore. compounds such as 2,5,6-triamino-4-pyrimidinone (II), 5-benzylamino-2,6-diamino-4-pyrimidinone, and pyrimidodiazepine III can act as substrates (54). The second reason why another tetrahydropterin may have been used as an electron donor is that BH_4_ is primarily used to combat oxidative stress (54), potentially limiting its supply during high temperature stress. Existing data demonstrate the likely involvement of P4H in the symbiotic relationship between *Hydra viridissima* and its photosymbiont *Chlorella* sp. A99 (55).

The second aspect we wish to consider is the impact of thermal stress on the coral reproductive cycle. Beyond the MAGI results regarding progesterone, analysis of existing metabolite ion counts from untargeted UHPLC-MS analysis of *M. capitata* (11) shows that predicted sex steroids in this species follow the expected increase in accumulation (e.g., testosterone, estrone, androstenedione) under ambient conditions during the mass spawning event (Figure 5). These metabolite levels are generally reduced under thermal stress, however, at T5, recovery to near ambient and wild sample levels are found for several compounds (e.g., estrone, androstenedione, testosterone; Figure 5). This suggests that *M. capitata* may be partially able to adapt to warming waters with regard to sex steroid production, although these preliminary results need validation. It should be noted that misregulation of coral spawning is an emerging problem in warming oceans (6) and was evident during our experiments when *M. capitata* spawning was highly attenuated in Kāne‘ohe Bay in June 2019 (D.B., H.M.P., unpublished data, June 2019). Our results demonstrate that thermal stress impacts the production of hormones linked to reproductive activity.

**Figure 5:**
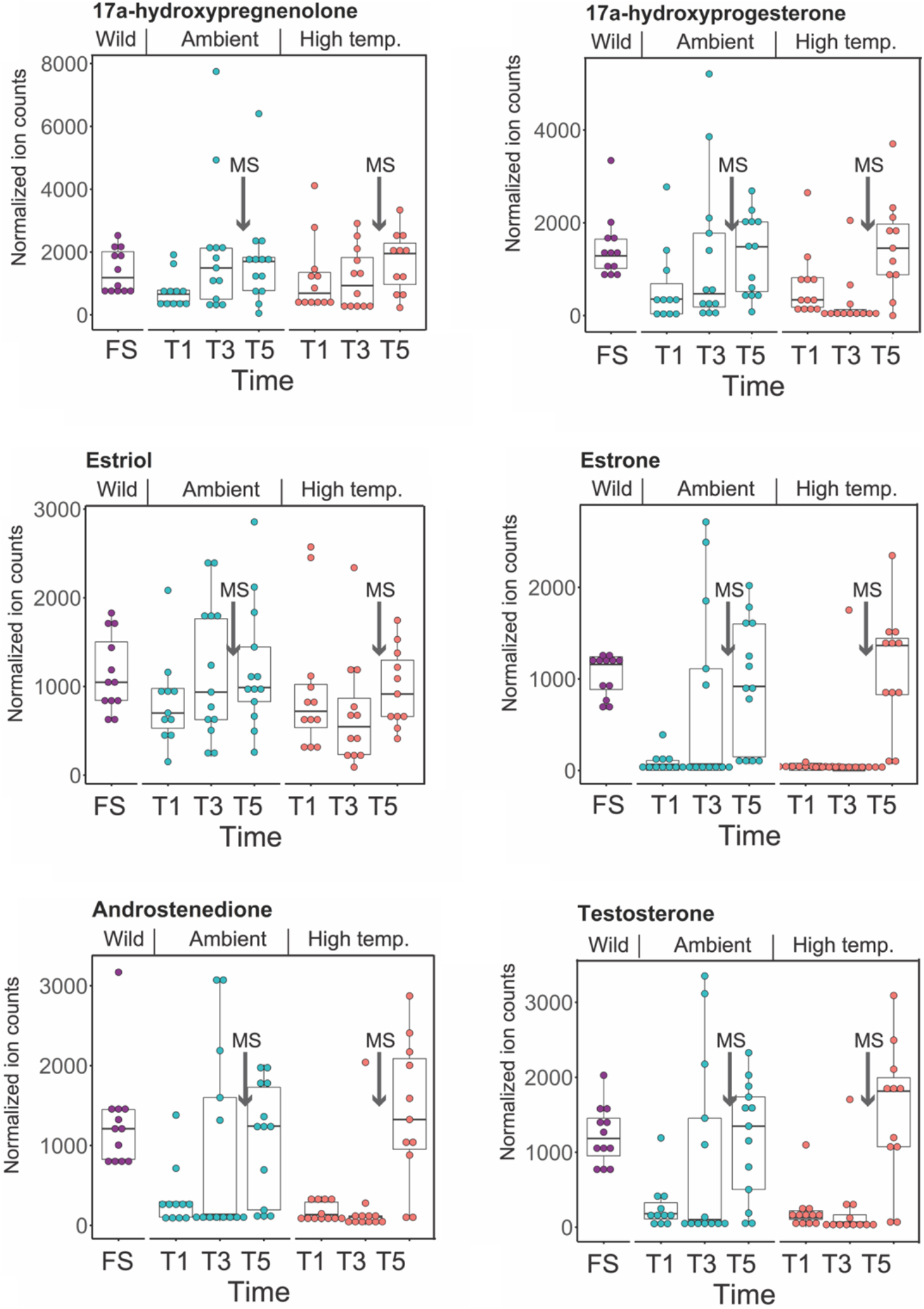
Analysis of sex steroids in *M. capitata*. Accumulation of a variety of predicted sex steroids in *M. capitata* nubbins over the duration of the ambient and thermal stress treatments as well as from wild populations collected after T5 from the same colonies used in the tank experiments [FS; for details, see (11)]. Each dot represents a single nubbin measurement. The pattern of metabolite accumulation suggests that these steroid levels increased at T5 (Figure 1B), which preceded the expected mass spawning event (arrow labeled with MS between T3 and T5) for this species in June 2019. The putative functions of these steroids are as follows: 17a-hydroxypregnenolone – a neuromodulator generated by the action of mitochondrial cytochrome P450 enzyme 17α-hydroxylase (CYP17A1) that is an intermediate in the delta-5 pathway of biosynthesis of gonadal steroid hormones and adrenal corticosteroids; 17a-hydroxyprogesterone – progestogen that is a chemical intermediate in the biosynthesis of androgens, estrogens, glucocorticoids, and mineralocorticoids; estriol – female sex hormone (weak estrogen), with a large amount produced in humans by the placenta; estrone – another female sex hormone (weak estrogen), binds to the estrogen response element and promotes transcription of genes involved in the female reproductive system functions; androstenedione - weak androgen steroid hormone, precursor of testosterone and other androgens; testosterone - primary male sex hormone involved in development of male reproductive tissues.

In conclusion, our results reveal the complexity of the thermal stress phenome in *M. capitata* that includes many genes involved in redox regulation, biomineralization, and reproduction. Significant effort will be needed to modify this polygenic trait in coral holobionts to boost resilience in the long term. Nonetheless, we have identified a number of novel genes that are promising candidates for functional analysis using the recently developed CRISPR/Cas9 tools for corals (20, 56, 57). It is important to remember that the algal symbionts of corals play a key role in holobiont biology and stress response vis-à-vis symbiotic nutrient cycling (58). Therefore, future gene-metabolite interaction analyses need to address *in situ* algal gene expression to address the integration of the host-symbiont response to thermal stress.

## Materials and Methods

### Metabolite data

All of the metabolomic data used here were produced as part of the Williams et al. (11) study and are available as the supplementary information associated with this manuscript (https://advances.sciencemag.org/content/suppl/2020/12/21/7.1.eabd4210.DC1).

### cDNA library preparation

The *M. capitata* samples used for transcriptomic analysis were a subset (3 replicates) of the same nubbins used to generate the metabolomic data in Williams et al. (11). Total RNA was extracted using liquid nitrogen and a mortar and pestle. RNA was isolated with the Qiagen AllPrep DNA/RNA/miRNA Universal Kit and strand specific cDNA libraries prepared using the TruSeq RNA Sample Preparation Kit v2 (Illumina) following the manufacturer’s instructions. This protocol includes poly-A selection to target eukaryotic cells, eliminating reads from the prokaryotic microbiome. Quality control for the libraries was done using an Agilent Bioanalyer, with library length being ~250 bp. Sequencing was performed on the NovaSeq (2×150bp) by the vendor GeneWiz. These RNA-seq data are available under NCBI BioProject ID: PRJNA694677 (see also Methods-table supplement 1).

### RNA-seq preprocessing

RNA-seq reads were trimmed using Trimmomatic v0.38 (mode ‘PE’; ILLUMINACLIP:adapters.fasta:2:30:10 SLIDINGWINDOW:4:5 LEADING:5 TRAILING:5 MINLEN:25) (59), only pairs for which both mates remained after trimming were used for subsequent analysis.

### Functional annotation

The reference *M. capitata* proteins were functionally annotated using the Uniprot database (release 2020_06). BLASTP (version 2.7.1+, parameters: *e*-value 1e^−5^ -seg yes -soft_masking true and pident ≥ 30%) was used to query the predicted proteins against the Uniprot (SwissProt + TrEMBL) protein database. Function assignment was based on the best hit criterion. Proteins without hits against Uniprot or annotated as “Unknown” were compared using BLASTP against the current NCBI nr database.

### Differential expressed genes (DEGs)

Expression of the available *M. capitata* genes (13) over the sequenced timepoints was quantified using Salmon v1.10 (--allowDovetail --validateMappings --seqBias ‒gcBias) (60). We kept only genes with a TPM value ≥ 5 in each sample. The R-package DESeq2 (61) was used to find the DEGs by comparing the ambient versus stressed condition for each time point.

### Co-expression networks

The R-package DGCA (62) was used to determine the correlation between pairs of genes respectively for each time point. The pairwise correlation was calculated with the function matCorr using Pearson method. The functions matCorSig and adjustPVals were used to calculate and adjust (with the Benjamini-Hochberg method) the correlation *p*-values, respectively. Only pairs with an adjusted *p*-value ≤ 0.05 were used to construct the co-expression networks. Module detection was done using the functions hclust (method = “average”) and cutreeDynamicTree (minModuleSize = 10 and deepSplit = TRUE).

### Differentially accumulated metabolites (DAMs)

We used the R-packge mixOmics (63) to detect differentially accumulated metabolites (DAMs) (VIP score ≥ 1 and FC ≥ 2). MAGI (10) was used with the default parameters to find the connection between DAMs and DEGs. MAGI results, using m/z and rt to define each metabolic feature, were filtered based on the MAGI scores (compound_score = 1, e_score_r2g ≥ 5, e_score_g2r ≥ 5, reciprocal_score = 2). We checked each reaction manually for DAMs and DEGs of interest.

Briefly, MAGI relies on a biochemical reaction network to numerically score the consensus between metabolite identification and gene annotation and generates metabolite−gene associations. Liquid chromatography-Mass Spectrometry (LCMS) features, which are defined by the combination of a unique retention time and mass-to-charge ratio (m/z) (64), and protein or gene sequence FASTA files serve as checks on each other. The likelihood of identifying an LCMS feature/gene function increases if there is a gene function/metabolite feature to substantiate that metabolite identity/gene function. The putative compound identification and input sequences are then connected to biochemical reactions by a chemical similarity network and evaluated based on sequence homology against a reference database (10). The MAGI score is a geometric mean of the homology score, reciprocal score, reaction connection score, and compound score, representing the probability and strength of the gene-metabolite association. The reaction connection score is not set by the user and shows how each metabolite is linked to genes and reactions. The compound score was set so that all metabolites would be given equal weight during data integration. The homology score, which is a composite of the reaction-to-gene (r2g) and gene-to-reaction (g2r) results, was set to 5, meaning that the reciprocal BLAST results are of the highest level. The reciprocal score, a direct representation of whether or not the r2g and g2r searches correspond to the same reaction, was set to 2, which requires reciprocal agreement.

## Supporting information

Supplemental File 1 cytoscape

## Acknowledgements

None.

## Supplements

**Figure 1-table supplement 1. Network size and gene expression direction** of individual modules for TP1, TP3, and TP5.

**Figure 4-table supplement 1. MAGI output at TP5, showing the highest scoring gene-metabolite interactions with a MAGI score ≥ 5.** The gene annotations, analyte identifications, MAGI scores, and reaction IDs are shown for both genes (GRT5) and metabolites (MRT5) at TP5. Rows highlighted in blue indicate redox reactions. Entries in the bold text take part in the same biochemical reaction.

**Methods-table supplement 1. Illumina RNA-seq data** generated from *Montipora capitata* as part of this study (NCBI BioProject ID: PRJNA694677).

**Supplementary File 1.** Cytoscape file containing the full networks and modules with gene and network information for the TP1, TP3, and TP5 gene co-expression results.

**Supplementary File 2.** Fasta files of *M. capitata* dark genes present in TP1 modules that are shown in Fig. 1C.

