## Supplemental Figures for "Multi-omic characterization of the thermal stress phenome in the stony coral *Montipora capitata*"

Title

Coral thermal stress, gene co-expression analysis, *Montipora capitata*, multi-omics, network analysis.

**Figure 1-figure supplement 1: Maximum likelihood (IQ-Tree) phylogenetic analysis of *M. capitata* dark gene g36545** done using default parameters and 1000 ultrafast bootstrap replicates. The results of the bootstrap analysis are shown on the branches when >60%. The legends show the expected substitution rate for the protein dataset.

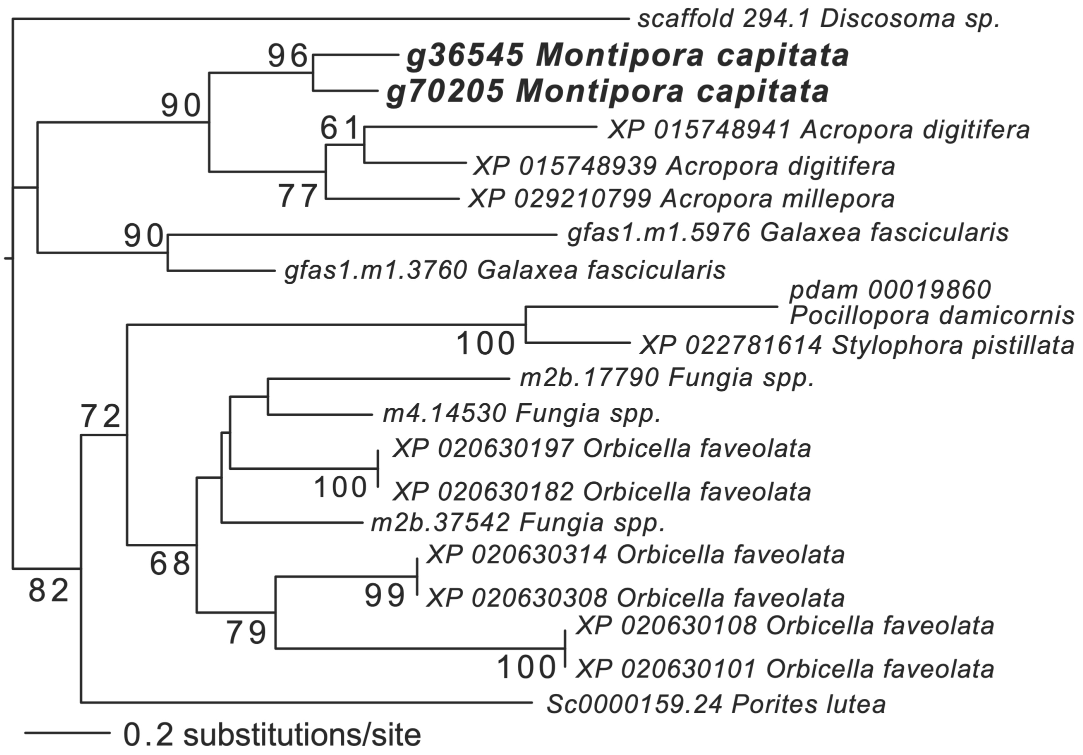

**Figure 2-figure supplement 1: *M. capitata* networks of differentially expressed genes at TP3 (top) and TP5 (bottom) showing the different gene modules and their interactions.** Red nodes are up-regulated, green nodes are down-regulated, and yellow nodes are dark genes. Edges that are purple (A) or blue (B) indicate downregulated genes, whereas edges that are orange (A) or red (B) indicate upregulated genes.

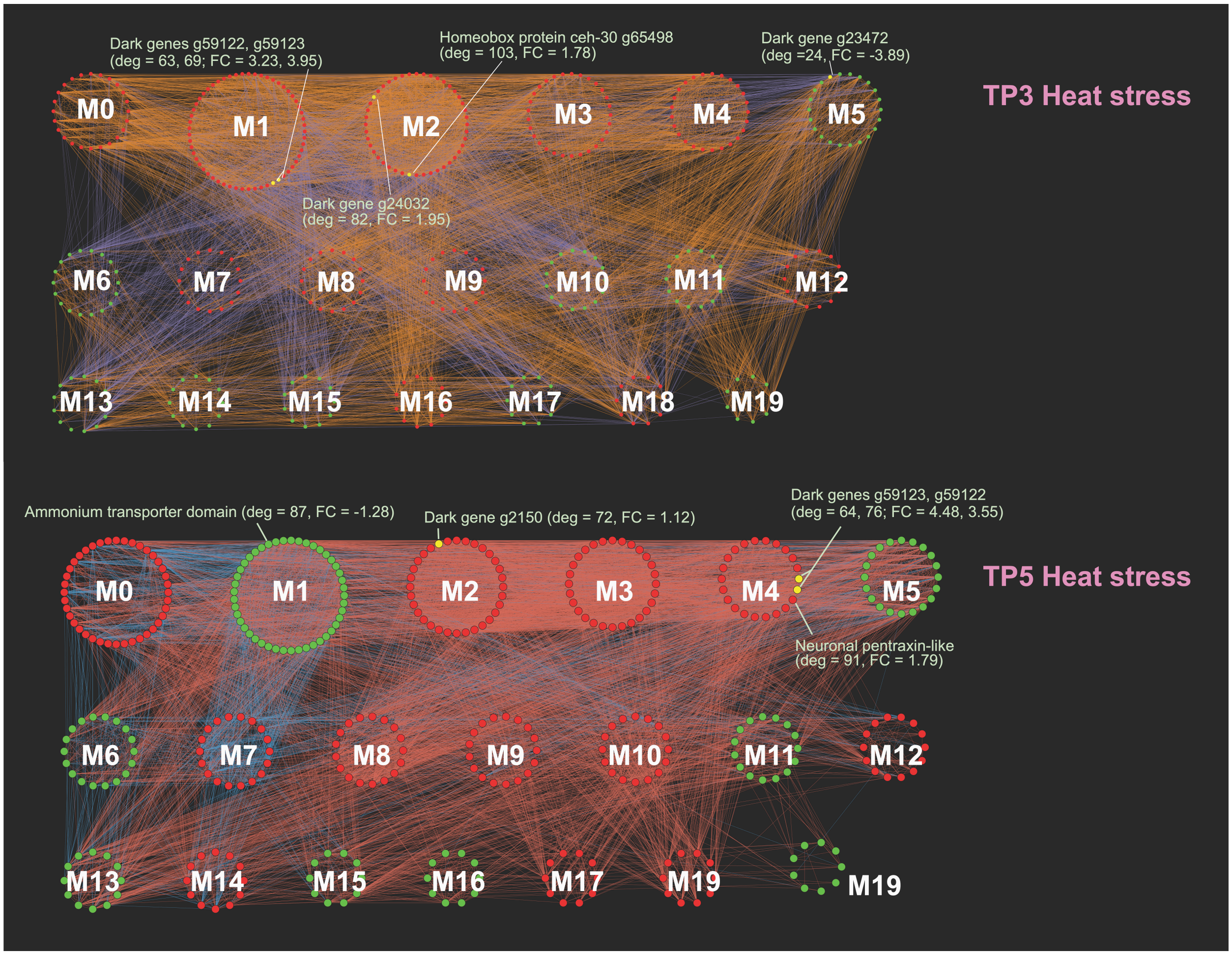

**Figure 2-figure supplement 2: Maximum likelihood (IQ-Tree) phylogenetic analysis of coral CARP5 homologs inferred using default parameters and 1000 ultrafast bootstrap replicates.** The results of the bootstrap analysis are shown on the branches when >60%. The legend shows the expected substitution rate for the protein dataset. Complex and robust coral species are shown in brown and blue text, respectively. Four putative CARP5 paralog clades are indicated. The thick branches mark a major gene duplication event in the common ancestor of complex and robust coral species.

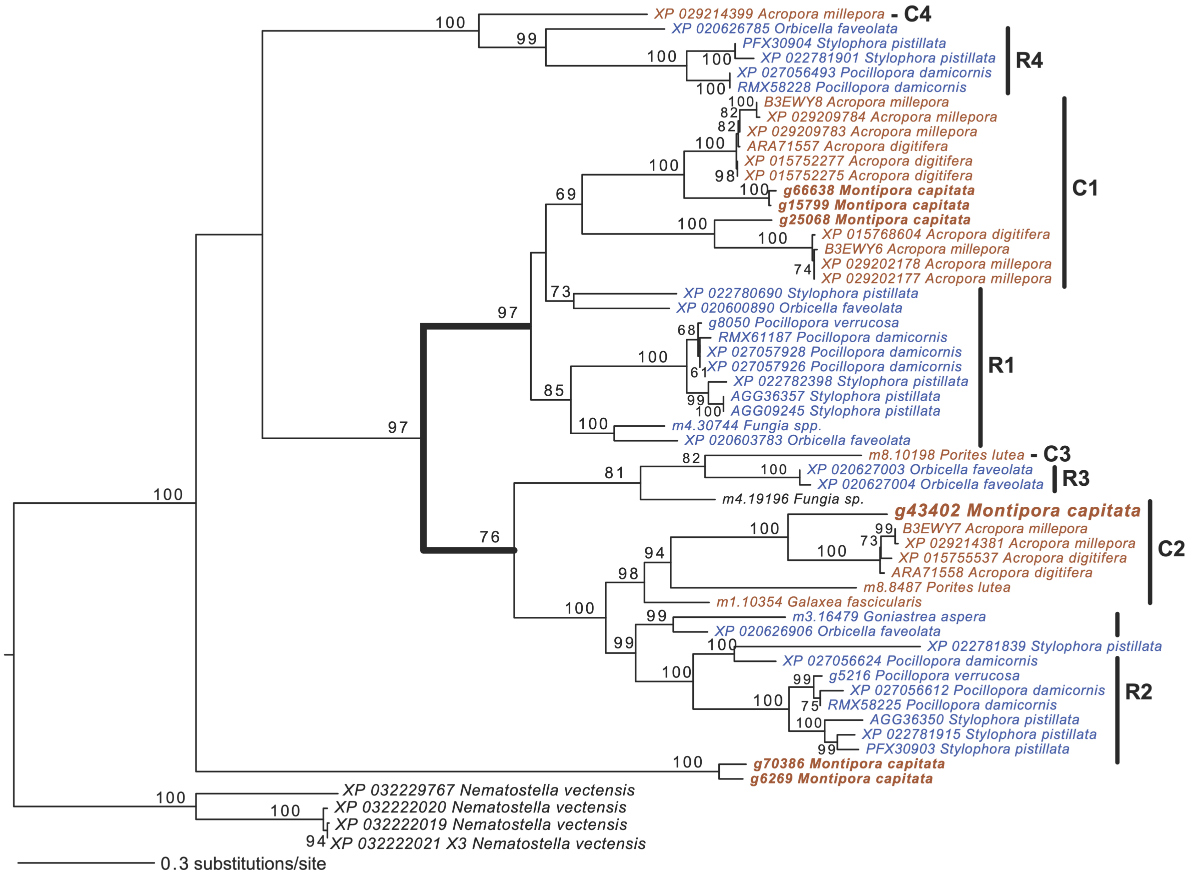
