## Supplementary material for "Multi-omic characterization of the thermal stress phenome in the stony coral *Montipora capitata*": Figure 4-table supplement 1

**Supplementary Table S2.** MAGI output at TP5, showing the highest scoring gene-metabolite interactions with a MAGI score $\geq$ 5. The gene annotations, analyte identifications, MAGI scores, and reaction IDs are shown for both genes (GRT5) and metabolites (MRT5) at TP5. Rows highlighted in blue indicate redox reactions. Entries in the bold text take part in the same biochemical reaction.

| Gene annotation | Compound name | InChI Key | MAGI score | GT5 | MRT5 | Reaction ID |
|---|---|---|---|---|---|---|
| Abhydrolase domain-containing protein 4 | Isobutyric acid | KQNPFQTWMSNSAP-UHFFFAOYSA-N | 5.129376 | UP | UP | RXN-668 |
| Alcohol dehydrogenase [NADP(+)] | 5alpha-Androstane-3beta,17beta-diol | CBMYJHIOYJEBSB-YSZCXEEOSA-N | 9.052214 | UP | DOWN | RHEA:16300 |
| Alcohol dehydrogenase [NADP(+)] | 2-Hexenal | MBDOYVRWFFCFHM-SNAWJCMRSA-N | 8.730280 | UP | NA | RXN-12299 |
| Alcohol dehydrogenase [NADP(+)] | Acrolein | HGINCPLSRVDWNT-UHFFFAOYSA-N | 8.676901 | UP | UP | RHEA:12170 |
| Alcohol dehydrogenase [NADP(+)] | (1R,2R)-Cyclohexa-3,5-diene-1,2-diol | YDRSQRPHLBEPTP-PHDIDXHHSA-N | 8.592966 | UP | UP | 1.3.1.20-RXN |
| Alcohol dehydrogenase [NADP(+)] | 1-Butanol | LRHPLDYGYMQRHN-UHFFFAOYSA-N | 8.548746 | UP | NA | ENZRXN-201-RXN |
| **Alcohol dehydrogenase [NADP(+)]** | **L-Gulonic acid** | **RGHNJXZEOKUKBD-QTBDOELSSA-N** | **7.707037** | **UP** | **UP** | **GLUCURONATE-REDUCTASE-RXN** |
| | ***aldehydo* -D-glucuronate** | **IAJILQKETJEXLJ-QTBDOELSSA-N** | **7.305923** | **UP** | **NA** | |
| Betaine--homocysteine S-methyltransferase 1 | (2R)-2-Ammonio-4-(methylthio)butanoate | FFEARJCKVFRZRR-SCSAIBSYSA-N | 10.873928 | NA | DOWN | RHEA:22339 |
| Cytochrome P450 10 | (25R)-3alpha,7alpha,12alpha-Trihydroxy-5beta-cholestan-26-al | USFJGINJGUIFSY-IUFSEJPUSA-N | 6.595443 | NA | NA | CHOLESTANETETRAOL-26-DEHYDROGENASE-RXN |
| Cytochrome P450 10 | (25R)-3alpha,7alpha-Dihydroxy-5beta-cholestan-26-oic acid | ITZYGDKGRKKBSN-RXDNHGQQSA-N | 6.595443 | UP | DOWN | RXN-9844 |
| Cytochrome P450 1A1 | 6-Hydroxymelatonin | OMYMRCXOJJZYKE-UHFFFAOYSA-N | 7.281551 | UP | UP | RXN-11056 |
| Cytochrome P450 1A1 | Progesterone | RJKFOVLPORLFTN-LEKSSAKUSA-N | 6.312111 | UP | UP | RXN66-355 |
| Cytochrome P450 3A24 | Eucalyptol | WEEGYLXZBRQIMU-UHFFFAOYSA-N | 8.436319 | UP | UP | RXN-13133 |
| Dimethylaniline monooxygenase [N-oxide-forming] | N,N-Dimethylaniline N-oxide | LKQUDAOAMBKKQW-UHFFFAOYSA-N | 11.203145 | UP | UP | RHEA:24470 |
| DLH domain-containing protein | 4-Carboxymethylenebut-2-en-4-olide | AYFXPGXAZMFWNH-UHFFFAOYSA-N | 5.990086 | UP | UP | RHEA:12373 |
| FAD-linked oxidase | (R)-6-Hydroxynicotine | ATRCOGLZUCICIV-SECBINFHSA-N | 6.846279 | UP | UP | RHEA:10015 |
| Glutamine amidotransferase type-1 domain-containing protein | cis-4-Hydroxy-D-proline | PMMYEEVYMWASQN-QWWZWVQMSA-N | 6.300669 | DOWN | NA | RXN-8003 |
| Glutamine amidotransferase type-1 domain-containing protein | 1-Aminocyclohexa-3,5-diene-1,2-diol | PWWIBLJDJCLRSC-UHFFFAOYSA-N | 6.051894 | UP | UP | RXN-15250 |
| Glutamine amidotransferase type-1 domain-containing protein | N(5)-Phenyl-L-glutamine | VMNRUJGOLBSEPK-VIFPVBQESA-N | 5.977851 | UP | UP | RXN-15251 |
| Guanylate cyclase (Fragment) | Pyrophosphoric acid | XPPKVPWEQAFLFU-UHFFFAOYSA-N | 10.689955 | DOWN | UP | RHEA:13665 |
| Inositol oxygenase | D-Glucuronic Acid | AEMOLEFTQBMNLQ-AQKNRBDQSA-N | 7.260806 | UP | NA | RHEA:23697 |
| L-tyrosine decarboxylase | Tyramine | DZGWFCGJZKJUFP-UHFFFAOYSA-N | 5.953216 | NA | UP | RHEA:14346 |
| 1-pyrroline-5-carboxylate synthase | D-Glutamic acid | WHUUTDBJXJRKMK-GSVOUGTGSA-N | 11.232582 | UP | UP | RHEA:24889 |
| Delta-1-pyrroline-5-carboxylate dehydrogenase, mitochondrial | 4-Hydroxyglutamate semialdehyde | XCXUZPXOFFRGGP-DMTCNVIQSA-N | 11.232582 | NA | NA | RXN-14472 |
| **Phe-4-hydroxylase** | **Tyrosine** | **OUYCCCASQSFEME-QMMMGPOBSA-N** | **11.494889** | **NA** | **NA** | **RHEA:20273** |
| | **4a-Hydroxytetrahydrobiopterin** | **KJKIEFUPAPPGBC-UHFFFAOYSA-N** | **10.909811** | **DOWN** | **UP** | |
| | **Phenylalanine** | **COLNVLDHVKWLRT-QMMMGPOBSA-N** | **10.804993** | **NA** | **NA** | |
| S-(hydroxymethyl)glutathione dehydrogenase | Acetaldehyde | IKHGUXGNUITLKF-UHFFFAOYSA-N | 10.629627 | UP | DOWN | RHEA:25290 |
| Glycerate dehydrogenase | D-Glyceric acid | RBNPOMFGQQGHHO-UWTATZPHSA-N | 9.974234 | DOWN | DOWN | RHEA:18658 |
