## Supplemental File 2 for "Multi-omic characterization of the thermal stress phenome in the stony coral *Montipora capitata*"

Module M0
>augustus.g59123.t1
MNSCSLFVAIVLLLHITKRGFSSPIPCKEGKFFEASVFDYFNCSVCEGQAHFTNCGLCCGSANATTTSSFSSTSSSDPTS
RPGSEKSESVNPCDLKFSFRENTLPIGIAFVTGFLAALVIVGIAKLVIRCRSKSKKRHPTELTPPQQETQPGLRASPMVI
GLDNGTGTNPNK*
>augustus.g46336.t1
MVPFRWLLVFAFLPMSSAFASITISQVPVLSSPVTVDAPVVVSSSSVQTFSVSLSNGTSTVSLSVSARAAVLMSSNSLGT
QSPSSSTIVTNRLPLTTHGTVSSSLTSRIISPSAAFTGTSTASTSVEATTPSVTDKTKSEFDWTWTVLGIVIGLIVIIST
MIVAGIILKRRGSAEFGKNVAHEMTAAQIENALASSNPVEVQT*

Module M1
>augustus.g695.t1
MGRFSNLTTKSSSSGDNLSLASCDSAFVVGHWLATGQSLLDYQRNNDVSSSNTRPYPIKMKTAIVFLALVMMTVGSVAEL
NARCDGQQPDIAEGRYFIENVETGRYLFQTGDKMKGNRGDEGGWLQAPEEVGSDANYYNRAYWKIIPQGDGKYFIENMET
QRYLFQTGDKMKGNRGDEGGWLQAPLRLDQMPTTTIEPTGR*
>augustus.g56482.t1
MSSRRRAVRVFQEESRFSTKQSRTETQGKKVTTMEKKLLCVVLIMLLMVSFNSAQPNQRIGKRNFQAVIKKREYCARARQ
ICSPTEDDVIYDGHMPDVYP*
>augustus.g4333.t1
MNPTFSVFAKENFKTESYRPVWQKSPTPLLDFKMTFKLTVACFIIASLMAVEAAKMPRTNHKHGIHEEARYRAHFLKDFL
DRRQEFLDLRSCVALRGYGCENNNAKCCREGNPYTGKMRKCVNVGSFSEP*
>augustus.g43083.t1
MSLIFVTMILCLSLLINFTGGQSQNWGCLFRPCRKRKLQAKSDSTSNEDTMNLEQRELNPKAKILLAHLMRRSYKRPQEF
RTDYADDDY*
>augustus.g49521.t1
MGEEETRMDPKCDNKTFATFYCPLFYKPAMYLFCKNARDPNNPAVAGKYFTEDYVLTRSDLPILENNLKMAQDIKTSIQK
QEEDAKKHLCTHFSKAGKLLPGHNHPKTFDEKLTGKGKVRARDYRIFNVPHPSHQPLLLSLVVESVEMNYESTEAHTFSK
TDGNNLKAGARVFAGNVTYSFLEVTVDTGLDVTERGATAVAEARGYIFQVLGVYADQTFARFFGFHVGPSVGFDLQSYVL
KGKVLGAEMKARTTLVEAQDGRYYVHLGAGFTTGAKVEDGAIQMKVLGSGFSVGRWNGFSLFGNEFAIHIRDILGSK*

Module M2
>augustus.g59122.t1
MKTSKYKRSFEIKQTMNYNFSLLALCIYFLWLFVGRSEGSVGSLCEDGKFFDQSKQQFISCHECNSEIQTCSLCCNQAES
DNVTTERSSNHHLQPLDTLGPDNKIMSSGNRDISPNVLVASIVGSVMLISLLTVILVILKRYAGRREEGTDLASGCGECL
RRPETLTVMTMENISESADTEGKFFEDSVFDYIDCSVCEGQAHFTNCKLCCGSANATPPSSFSSTSISSSDRTSLPGSEK
SESDVNPCDLKFSFRENTLPIGIAFVTGFLAALVIVGIAKLVIRCRSKSKKQHPTGYTPPQQETQPDLHGASPMVPGLTV
ELPQMTQELI*
>augustus.g36545.t1
MRITDLPRETFDEVCSVLGKDTTINWKKLMTESYRSLYLPKHVEEIGQKRFPADALLNDLADREVPLDDLLRELQKIGNK
KAVSIINKAIKRKDFNGGPPTQTITREPEESTATPSDRGQTETLPPYSIEESSTHGGDPCYPKLEEFMERISMLFPCMNP
LIWRTAH*

Module M3
>augustus.g13.t1
MSLLISLFVGSFLFCVGLEARIEDTVTVSAKNSNFSVSEINRSKLKSVFEKNGYKITEIQEKSRVNSGKEFNDTVIILST
PKAPCSEAPKEISAMKAQLLKAVKVQESVDKISALFNHLQADLNKQNTNGRLCQTSVKLLFKDALHKAVGYERPKRFLWG
RRRRRRRTSTTTTTTTTKTTTLTVTTTTTTTTTSSSG*
>augustus.g25275.t1
MGYTRVLLMAIIVAMITEKAWGFISYDVQSTIVGLMDQRDFTALQKVRLTQDNVRPIFRKDISGKAPPYYEISAGGHYYV
FSASSRTGDHRLVESGKGIRPTELLNKQALIKGETCDKYYRITAVSGLYTCENSKGLLVASTYDFLSDAPVDNVNLKNKT
YYYHLQTSLRLNLPEFNKEWQDLHKETKDRAEWTTEQILILTEKKQDASNTRYAEEALRPGSSVKVALGNRFSRVSIRFT
GTGRNTSGDDEPLEEWQRRLCQVLDRVCLNGSKKINKEAIEVAPNGISYIKLSVSEDKDRVRKILQNAEVGFMIEFHVSE
DGRHLLVRKRFAIDLSPSRKRRWSTWITFSVPEEGTFPDYKQHECCGCKSGCGPVAWAQIFAYYDRLAHASSRYGYSKDL
YRCKSGIAGSSACKAPLRLNTASVKKYVEHIRSQVDTFCLLKGGFTAHWNMKDSKLKSFYRSRQSGGTIYGYTSWLTSLP
GIYSSRIRDRAIEAVKQGYPAIVGIWSGLSQHYAVATKYRYRSQKNRICCGRWKTRYVHEFYYHMGWGGNHTASWRTAKA
FSAFVAKK*
>augustus.g30126.t1
MAVLVRSCCCGCKLKTGVLLLAIFSVIGGGYGIYSSFDKASKESSSPVFSKYSDVITNLLKTNGVFNIIVLLTSILLLGA
NILKNRFLVLPYFAWHVILLGYRLGVSIFFTVVWKGPFVYAVVIFSAISWLLSIYFLIVVYSFHEALREDPSGATAGYGP
ANPSGQAASRPPPPAVSHKSANFV*

Module M4
>augustus.g54458.t1
MFIWLLWMNPIRYKRGQGRAAPRKDSLKSAIFRSVCRSLKEVIIMKQTLVIVLVVFLVLSFINESFLWSLRQRRFLSRPV
VIHAMDNDAGAGPASLLAAREADDGVRVLSPETAFGEIPRVSAASRKDSLKSAIFRSVYRSLIEVIIMKQTLVIVLVVFL
VLSSINESDAWRRRRAGLFHKRNHNDKTHTIKTSQDDEGDKTKKHRFHDDKDTYEDADLL*
>augustus.g6815.t1
MTMNQRNNDVSSSNTRPYPIKMKTAIVFLALVMMTVGSVAELNARCDGQQPDIAEGRYFIENVETGRYLFQTGDKIKGDR
GDEGGWLQAPPEVGSDANYYNRAYWKIIPQGDGKYFIENQETQRYLFQTGEKIKGNRGDEGGWLQAPAEVGSDANYYNRA
YWKIIPQGDGKYFIENLETQRYLFQTGEKIKGDRGDEGGWLQAPPVVGSDANYYNRAYWKLLEQ*

Module M5
>augustus.g22745.t1
MNSFILILVPLMVFAFPNALAITCYKCTPAQNPLRTCTKPEELTSMECPTPNGNMSMNVSGMNMSMNVTGVNISLTYDAC
LTTRIMGNVPILGQLTSYYLSCGMQDPGSSSSMPNCSMTGSSICERAEQQVSGNGITIVSCDNTCCTTDNCNEIEQPSTT
PSNNEETTMASTTSGMDQVRPQFFGLLIVFAGILFRKIDQPF*
>augustus.g22744.t1
MCTPAQNPLRTCTKPEELTSMECPTPNGNMSMNVSGMNMSMNVTGVNISLTYDACLTTRIMGNVPILGQLTSYYLSCGMQ
NNGSSGSMPNCSMSQSGICKVAEEALIGTGITIVSCDNTCCTTDNCNEIEQPSTTPSNNEETTMASTTSGMDQVRPQFLG
LLMMFAGILFRKIDQPF*
>augustus.g31735.t1
MSENNNKDNPEIKITSNEIPYPPFKGRRNSEARGSRLVHKNGKHHVRSSNIPQRRERFLADFFTSFIDAKWRWVVVLYSA
GFLFSWCLFGTVYFIIFELRQKYDNGVLCVEKVDSWTSAFLFSVESQTTIGYGGRQITPECPEAIIFLLLQSLTGFMLST
SLLGLIFAKLSRPRPRAQTVMFSKHAVIAKQDGLLSLMFRVGDARKSQLLDVTVSLHCIQFRTTSTGQEILVSQQELSIC
TEHGIEVGQKIYPFLLLPLTVVHVIDERSPLYELGAQDLKFSRLELVAVLEGVVESTSMVTQARASYLAEEIVWGERFNP
ISVLRNIERGWKADFSSFDRTHKVGTPTLSAKEQRESKNKEDEKRKESLQISECQVWIQREEDTEFAPEKLEMHSKSNCK
AV*

Module M6
None
